# Enrichment of SARS-CoV-2 entry factors and interacting intracellular genes in peripheral immune cells

**DOI:** 10.1101/2021.03.29.437515

**Authors:** Abhinandan Devaprasad, Aridaman Pandit

## Abstract

SARS-CoV-2 uses ACE2 and TMPRSS2 to gain entry into the cell. However, recent studies have shown that SARS-CoV-2 may use additional host factors that are required for the viral lifecycle. Here we used publicly available datasets, CoV associated genes and machine learning algorithms to explore the SARS-CoV-2 interaction landscape in different tissues. We find that in general a small fraction of cells expresses ACE2 in the different tissues including nasal, bronchi and lungs. We show that a small fraction of immune cells (including T-cells, macrophages, dendritic cells) found in tissues also express ACE2. We show that healthy circulating immune cells do not express ACE2 and TMPRSS2. However, a small fraction of circulating immune cells (including dendritic cells, monocytes, T-cells) in the PBMC of COVID-19 patients express ACE2 and TMPRSS2. Additionally, we found that a large spectrum of cells (in circulation and periphery) in both healthy and COVID-19 positive patients were significantly enriched for SARS-CoV-2 factors. Thus, we propose that further research is needed to explore if SARS-CoV-2 can directly infect peripheral immune cells to better understand the virus’ mechanism of action.

## 1 Introduction

Severe acute respiratory syndrome coronavirus 2 (SARS-CoV-2) is a novel virus from the Coronaviridae family that has infected more than 100 million individuals, and has caused a rapidly unfolding global pandemic. A large number of infected individuals present no or mild symptoms and yet can spread the virus to others. However, some infected individuals develop severe acute respiratory distress syndrome resulting in coronavirus disease (COVID-19). This leads to a unique dysregulation of the immune system that is accompanied by a strong inflammatory response, cytokine storm and ultimately respiratory distress and viral sepsis (H. Li et al. 2020). However, the reason why the immune system enters such dysregulation is not yet clear. As a result, the exponential growth of the infections and failure to contain the infections has brought the world to a standstill. Thus, there is an urgent need to understand the mechanisms of infection and pathophysiology of SARS-CoV-2.

SARS-CoV-2 belongs to Betacoronavirus genera of Coronaviridae family and two other Betacoronaviruses (SARS-CoV and MERS-CoV) are known to infect humans. SARS-CoV-2 differs from the other two Betacoronaviruses; results in milder clinical manifestation in most individuals but has high transmission rate between humans (Petersen et al. 2020). SARS-CoV-2 uses spike protein S to infect human cells. The S protein of SARS-CoV-2 is homologous to the spike protein of SARS-CoV (>75% sequence identity) (Wu et al. 2020). The S protein of SARS-CoV has been shown to bind to the human ACE2 receptor for cell entry (W. Li et al. 2003) and requires target cell protease TMPRSS2 for its proteolytic priming (Matsuyama et al. 2010; Glowacka et al. 2011; Shulla et al. 2011). Due to the high sequence similarity in spike proteins of the two SARS coronaviruses, SARS-CoV-2 was postulated to use ACE2 as the entry receptor (Wu et al. 2020). Multiple studies have now confirmed that S protein of SARS-CoV-2 binds to human ACE2 for cellular entry and TMPRSS2 aids in its proteolytic activation (Hoffmann et al. 2020; Walls et al. 2020; Yan et al. 2020). Contrary to SARS-CoV, FURIN, a proprotein convertase was found to activate the S protein of SARS-CoV-2 by cleaving at the FURIN cleavage site found between the S1/S2 domains of the S protein. FURIN dependent activation along with TMPRSS2 and lysosomal cathepsins had cumulative effects on viral entry (Shang et al. 2020).

The presence of these entry factors specifically ACE2 and TMPRSS2 has been detected in bronchial & nasal epithelium and alveolar epithelial type II cells of the respiratory tract and in cells from ileum and colon of the digestive tract (Bertram et al. 2012; Hou et al. 2020; Zhao et al. 2020; Zou et al. 2020; Qi et al. 2020; Zhang et al. 2020). Hou et al. reported a decreasing gradient in gene expression of ACE2 and infectivity of SARS-CoV-2 from the proximal to distal respiratory tract, with the ciliated airway and AT2 cells being the main target for infection. Pathological changes due to SARS-CoV-2 have been reported to be found in lung, kidney, heart, blood vessels, liver, colon, and digestive tract (Yao et al. 2020; H. Li et al. 2020; W. Wang et al. 2020). Using single cell RNA-Seq datasets of 13 human tissues, Qi et al. reported expression of ACE2 by lung, liver, stomach, ileum, kidney and colon. Qi et al. postulated ANPEP, DPP4 and ENPEP as candidate co-receptors as they were coexpressed with ACE2 in the tissue (Qi et al. 2020). Sungnak et al. analysed multiple single cell RNA-Seq datasets of 17 different tissues and found ACE2 expression limited to airways, cornea, esophagus, ileum, colon, liver, gallbladder, heart, kidney and testis (Sungnak et al. 2020). While TMPRSS2 was expressed by a broader number of cell types and tissues, the cells coexpressing ACE2 and TMPRSS2 were found to be from the respiratory tree, cornea, esophagus, ileum, colon, gallbladder and common bile duct. However, several innate immune genes such as IDO1, IRAK3, NOS2, TNFSF10, OAS1 and MX1 were found to be coexpressed with ACE2+ respiratory epithelial cells. In lung and bronchial tissue datasets, FURIN was found to be expressed in bronchial transient secretory cells and was found to be expressed in both ACE2+/TMPRSS2+ and ACE2+/TMPRSS2-cells, thus reducing the proteolytic dependence of TMPRSS2 (Lukassen et al. 2020).

Although most studies have focused on ACE2, TMPRSS2 and FURIN. We still don’t fully understand all the mechanisms by which SARS-CoV-2 infects and interacts with human cells. Recently, using cell lines and mice experiments, CD147-spike protein was found to be an alternative route of viral entry into host cells via endocytosis (K. Wang et al. 2020). However, this was later shown to be incorrect with no evidence of CD147 being involved as the binding receptor for SARS-CoV-2 spike protein (Shilts et al. 2021). Two seminal studies, Zhou et al. and Gordon et al., have looked into genes associated with SARS-CoV-2 to gain further insights into the mechanisms of action of the virus and find potential (intra-)cellular targets for future therapies (Gordon et al. 2020; Zhou et al. 2020). Zhou et al. took a network medicine approach to repurpose drugs for SARS-CoV-2 and by studying the host factors that have been associated with previously known coronaviruses (4 human coronaviruses: SARS-CoV, MERS-CoV, HCoV-229E, and HCoV-NL63; 1 murine coronavirus and 1 avian coronaviruses) (Zhou et al. 2020). Gordon et al. cloned and expressed 29 SARS-CoV-2 proteins in human cells and affinity purification mass spectrometry to identify 332 high-confidence human proteins that interact with SARS-CoV-2 proteins (Gordon et al. 2020). Studying human SARS-CoV-2 specific protein-protein interactions aims to provide insights into host factors that may influence infection dynamics and provide potential therapeutic targets.

Despite the growing literature into SARS-CoV-2 infection, it is still not clear which all cell types can get infected by the virus. Most studies have considered the expression of ACE2 within a cell type as the determining factor of SARS-CoV-2 susceptibility. However, it is still not clear if the human cells that express ACE2 also express the virus associated genes in them. Additionally, it is now clear that immune cells play a crucial role in the infection dynamics of SARS-CoV-2 and the underlying dysregulation of the immune system is distinct from other coronaviruses (Blanco-Melo et al. 2020). Several preprint studies have shown that immune cells can be infected by SARS-CoV-2. One such study showed that immune cells (CD4+ T-cells, CD8+ T-cells, B-cells and monocytes) in peripheral blood mononuclear cells (PBMCs) and lung tissue were infected with SARS-CoV-2 and infection was independent of ACE2 expression (Pontelli et al. 2020). Another study found that SARS-CoV-2 infects circulating monocytes and macrophages that further drive immunoparalysis and advance disease progression (Boumaza et al. 2020). ACE2 expressing tissue resident CD169+ macrophages were found to be infected by SARS-CoV-2 in the spleen and lymph node, that further progressed neutralization of the tissue (Feng et al. 2020). Such studies are essential to shed light on the role of immune cells in the disease progression. With the limited number of such studies focusing on fewer immune cells, it is still not clear which all immune cells in the different tissues and periphery are susceptible to infection and can potentially support the viral infection. To address this problem, we built upon Sungnak et al. to not only explore ACE2 and TMPRSS2 expression but also the expression of SARS-CoV-2 associated host factors in several tissues and cell types including lung, kidney and immune cells using publicly available single cell RNA-Seq data. We additionally extended this approach to study circulating immune cells curated from bulk RNA-Seq datasets. And, also included single cell RNA-Seq dataset of PBMCs from COVID-19 patients of varying severity, to explore the expression of SARS-CoV-2 associated host factors in infected cells.

## 2 Methods

### 2.1 SARS-CoV-2 associated gene sets

The Zhou gene list consisted of 119 genes involved in the virus-host protein interactions of several HCoVs, such as SARS-CoV, MERS-CoV, IBV, MHV, HCoV-229E, and HCoV-NL63 (Zhou et al. 2020). The Gordon gene list consisted of 332 genes that are physically associated with 26 of the SARS-CoV-2 proteins identified using affinity-purification mass spectrometry (Gordon et al. 2020). The 28-EF gene list consists of SARS-CoV-2 and coronavirus-associated receptors and entry factors (Singh, Bansal, and Feschotte 2020). The Integrin gene list comprises all known human Integrins (Hynes 2002).

### 2.2 Single cell data analysis

The normalised and preprocessed single cell RNA-Seq datasets were acquired from the covid19 cell atlas (www.covid19cellatlas.org). The normalisation, preprocess steps and cell type annotation of all the single cell RNA-Seq datasets performed using the scanpy suite are described by (Sungnak et al. 2020). The total fraction (Figure 1a) was calculated in percentage for all cells in a dataset for all datasets. The fraction (Figure 1b-g) was calculated in percentage for each annotated cell type for all datasets. The gene score (Figure 2b-g) for each of the above gene lists and for all cell types in all datasets was calculated using the ‘tl.score_gene’ function from the scanpy suite (Wolf, Angerer, and Theis 2018). The gene score is the difference in the average expression of the gene list to the average expression of background (all other) genes. This score reproduces the gene score in Seurat pipeline (Satija et al. 2015). Positive gene scores show that the cell expresses the given set of genes higher than the background. Wilcoxon non-parametric statistical analysis was performed with the alternative set to ‘> 0’ on the gene scores for every cell type in all datasets to measure significance of gene score. P values were binned in ranges of values ⩽ 0.001, ⩽ 0.01, ⩽ 0.05 and NS (non-significant, when p value > 0.05). As a control we computed gene score and performed Wilcoxon non-parametric statistical test using randomly selected genes. We did not find significant enrichment of randomly selected genes in majority of tissues and cell types (data not shown).

**Figure 1:**
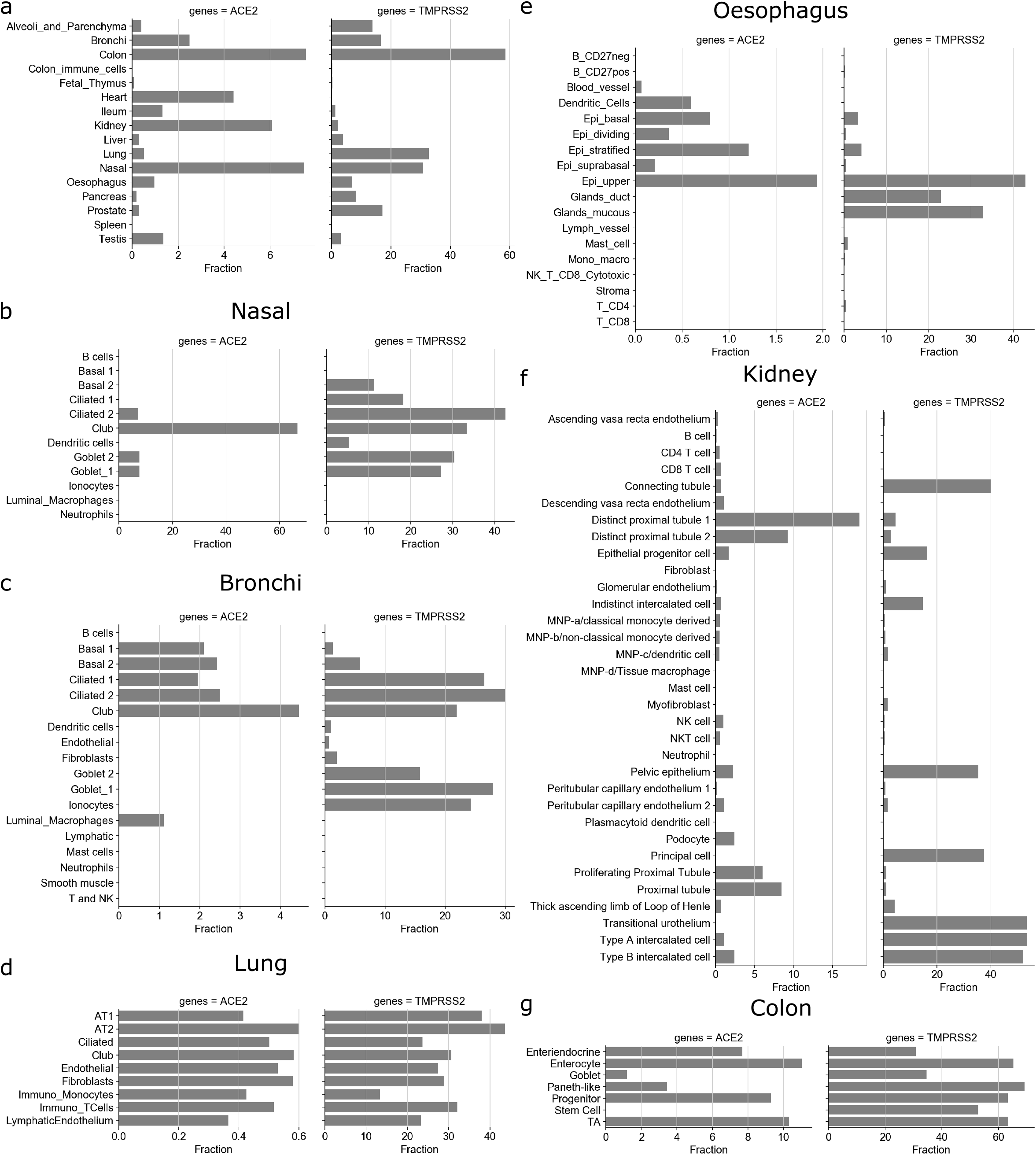
Fraction of ACE2 and TMPRSS2 expressing cells. a. Fraction of cells (as percentage in y-axis) that express ACE2 and TMPRSS2 in different tissues. Fraction of cells within tissue type: b. Nasal, c. Bronchi, d. lung, e. Oesophagus, f. Kidney, and g. colon that expresses ACE2 and TMPRS22.

**Figure 2:**
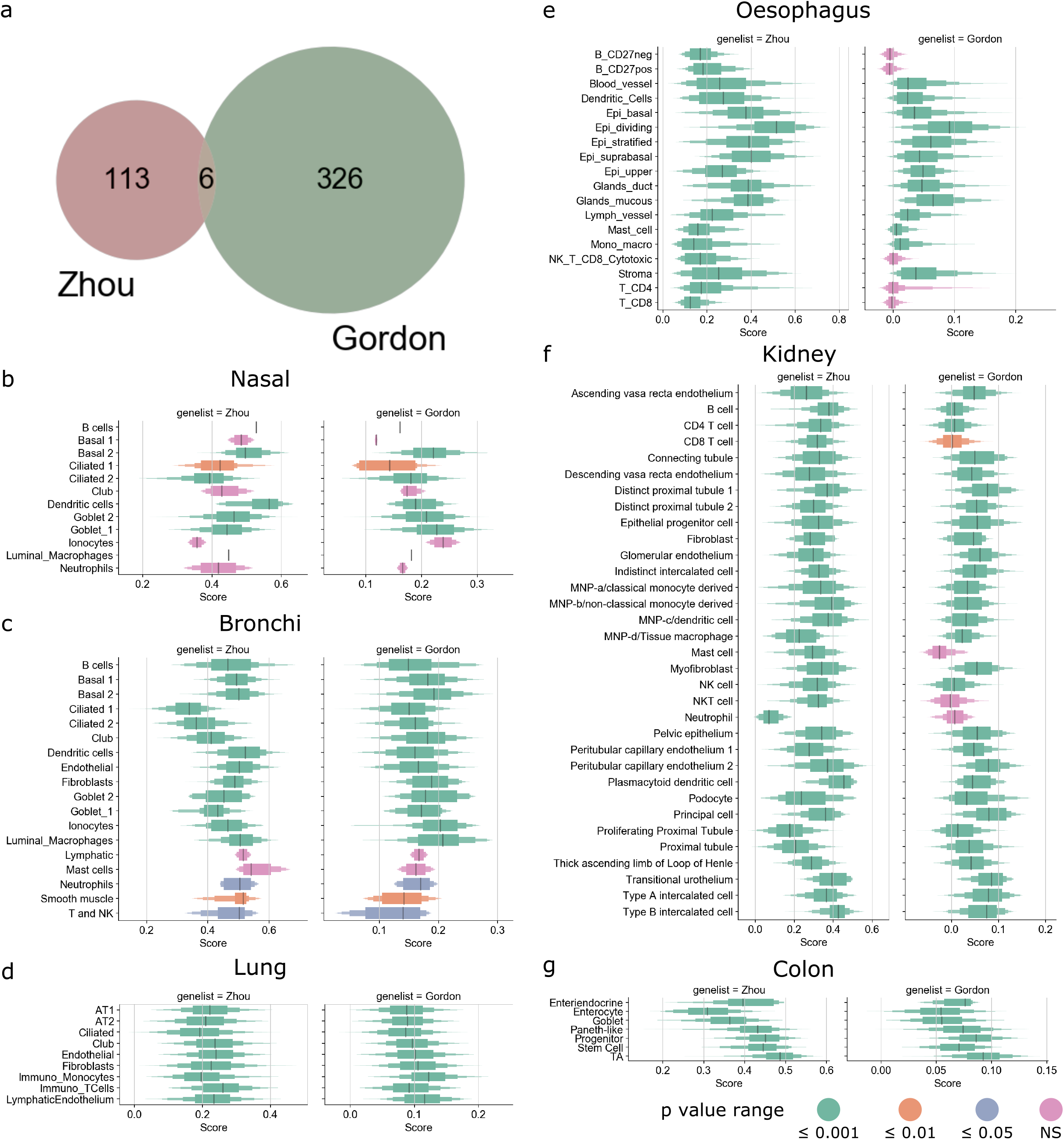
SARS-CoV-2 host factor genes. a. Venn diagram showing overlap of SARS-CoV-2 host factor genes between the Zhou and Gordon gene lists. Boxplot showing the distribution of gene score of Zhou and Gordon genes for different cell types: b. Nasal, c. Bronchi, d. lung, e. Oesophagus, f. Kidney, and g. colon. Black line represents median, height of box corresponds to number of cells in score range. Color of the box corresponds to the Wilcoxon p-value range (see legend) computed with the alternative set to > 0.

### 2.3 DIME on Immunome (bulk RNA-Seq)

The DIME tool (Devaprasad, Radstake, and Pandit 2019) identifies the top gene (from an input gene list) and top cell type cluster within an expression dataset by using non-negative matrix factorization (NMF). The shiny app implementation of DIME tool is available on bitbucket for installation and use (https://bitbucket.org/systemsimmunology/dime/src/master/). The DIME was applied on the immunome dataset available as default expression dataset in the tool. The immunome dataset comprises bulk RNA-Seq gene expression data of 27 immune cells of which 11 are myeloid and 16 are lymphoid. All datasets used in the construction of the immunome are from publicly available datasets (Devaprasad, Radstake, and Pandit 2019). The cells used here are from unstimulated (except for macrophages, that were monocyte derived) healthy donors. The DIME was run on the immunome using the Zhou, Gordon, 28-EF and Integrin gene lists to identify key cell types important for these gene lists (Figure 3). The highest ranking cluster was identified using Frobenius norm (Devaprasad, Radstake, and Pandit 2019). The top 25 genes for each ranking cluster are displayed (Figure 3). Reactome pathway enrichment analysis was performed on the top 25 percentile genes in each ranking cluster for the DIME results of the different gene lists (Supplementary Figure 5).

**Figure 3:**
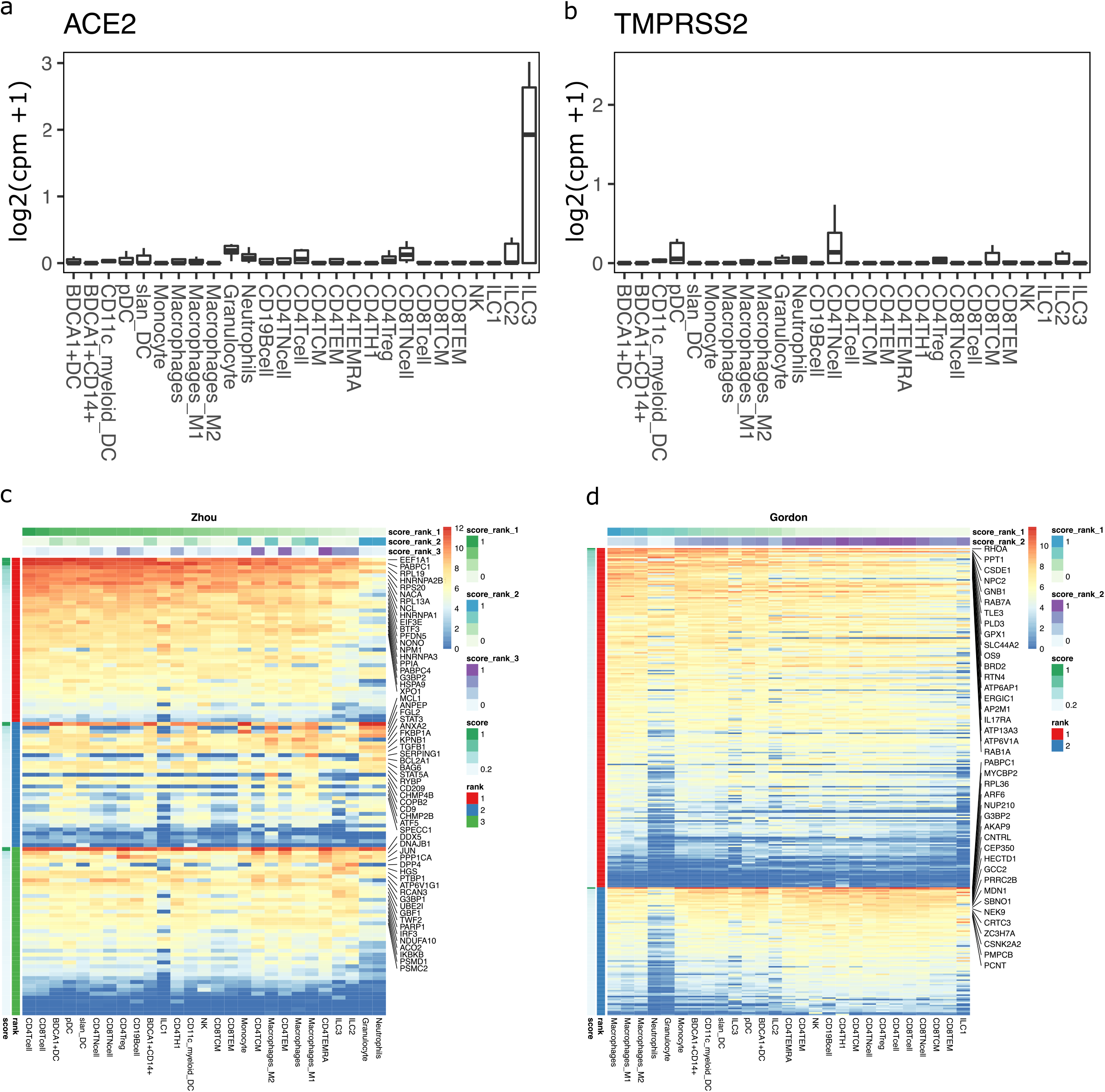
DIME enrichment of SARS-CoV-2 host factors in circulating immune cells. Expression of a. ACE2 and b. TMPRSS2 in circulating immune cells from the bulk dataset. Expression values are in log2(cpm+1). DIME Heatmap showing ranked enrichment of c. Zhou, and d. Gordon gene list in the circulating immune cells. The ranks depict the clusters as identified by DIME (see methods). Top 20 genes for each rank are shown. The cells are ordered based on the score of the rank 1 (Top weighted rank). Expression values are in log2(cpm+1).

### 2.4 Protein expression of the SARS-CoV-2 associated genes in circulating immune cells and tissues

To check the protein expression of the key SARS-CoV-2 associated genes identified by DIME (Figure 3 and Supplementary Figure 4), we used two publicly available protein expression datasets, namely, immprot and the human protein atlas (HPA). The immprot dataset comprises protein expression of 28 circulating primary human hematopoietic cell populations from healthy donors identified by high-resolution mass spectrometry-based proteomics (Rieckmann et al. 2017). The HPA dataset comprises protein expression of peripheral cells (mostly non-immune cell types) from 32 tissues (Uhlén et al. 2015). We checked the protein expression of the top 25 percentile genes of each cluster from the DIME results of the SARS-CoV-2 associated gene lists in the two protein expression datasets. Additionally, we also computed the Pearson correlation between the RNA-Seq and protein datasets for the cells that were common between the two datasets.

### 2.5 Plotting and visualization

DIME was implemented and visualized in R (version = 3.6.1). The single cell RNA-Seq analysis was performed in python (version = 3.6) using scanpy package and Wilcoxon statistics using stat package from python.

## 3 Results

### 3.1 ACE2 expressing cells across different single cell RNA-Seq

We first assessed the expression of ACE2 and TMPRSS2 in different tissues and cell types using 16 single cell RNA-Seq datasets. The ACE2 gene is expressed only in a small fraction of cells in different tissues. Among which colon (7.55 %), nasal (7.47%), kidney (6%), heart (4.4%), bronchi (2.49%), testis (1.35%) and ileum (1.33%) have the highest fraction of ACE2 expressing cells (Figure 1a). Despite being one of the most affected tissues, only 0.52% of cells expressed ACE2 in the lung. Interestingly, TMPRSS2 is expressed in many more cells in different tissues: colon (58.58%), lung (32.83%), nasal (30.87%), prostate (17.1%), bronchi (16.53%), alveoli and parenchyma (13.8%). Interestingly, ACE2 gene was expressed in less than 0.01% of spleen and in colon immune cells (Figure 1a).

We further studied the expression of ACE2 and TMPRSS2 in different cell types in each of the tissues (Figure 1b-g, Supplementary Figure 1). In corroboration with Hou et al., the decreasing gradient of ACE2 expression was observed in the respiratory tract from nasal to bronchi to lung to alveoli and parenchyma (Figure 1a-d). The club cells had the highest fraction of ACE2 expressing cells in nasal (66.6%) and bronchi (4.46%). Interestingly, the fraction of TMPRSS2 expressing ciliated-1 cells showed an increasing trend from nasal (18.18%) to bronchi (26.47%). We found that for lungs only a small fraction (0.36%-0.6%) of cells for each cell type expressed ACE2 gene (Figure 1d). For example, 0.41% of AT1 (alveolar type I) cells and 0.6% of AT2 (alveolar type II) cells expressed ACE2 gene (Figure 1d) while more than one-third of AT1 (38%) and AT2 cells (43.68%) expressed TMPRSS2. The low ACE2 expression in the lungs has also been reported in other studies (Zou et al. 2020; Qi et al. 2020).

Other organs that are known to undergo pathological changes due to SARS-CoV-2 include the kidney, heart, vessels, liver, colon, and digestive tract (Yao et al. 2020; H. Li et al. 2020; W. Wang et al. 2020; Lindner et al. 2020). In Oesophagus, the part of the upper gastrointestinal (GI) tract of the digestive system, the fraction of cells expressing ACE2 include 1.93% of the upper and 1.21% of stratified epithelial cells (Figure 1e). Colon had the highest fraction of ACE2 expressing cells, from the progenitors (9.3%) and transit-amplifying (TA) cells (10.31%) to the differentiated enterocytes (11%) expressing ACE2 (Figure 1g). Overall, the ACE2 expressing cells (Figure 1a) were lower in the oesophagus (0.97%) compared to tissues of the lower GI tract such as ileum (1.33%) and colon (7.55%). ACE2 was expressed at higher frequency in renal tubular cells such as distinct proximal tubule 1 (18.46% cells), distinct proximal tubule 2 (9.27% cells), and proximal tubule (8.45% cells), and SARS-CoV-2 has been shown to damage kidney in patients (Figure 1f) (Yao et al. 2020). Similarly, small fraction of cells in each cell type found in heart (<18% stromal cells, 3.26% adipocytes, etc), colon (7.7% of enteroendocrine, 1.17% goblet cells, etc), and ileum (0.24% endothelium, 0.86% goblets, etc.) expressed ACE2 gene (Supplementary Figure 1). Interestingly, for same/similar cell types found in different peripheral organs, we found different fractions of them to be positive for ACE2 expression. For example, 13% of fibroblasts expressed ACE2 gene in heart while only 0.2% and 0.17% of fibroblasts expressed ACE2 gene in alveoli and ileum, respectively (Supplementary Figure 1). Similarly, 41.65% of enterocytes expressed ACE2 gene in ileum while 11% of enterocytes expressed ACE2 gene in colon (Figure 1, Supplementary Figure 1). This potentially indicates that cells with the same/similar phenotype might vary in terms of ACE2 expression based on the local tissue environment.

### 3.2 Enrichment of SARS-CoV-2 associated genes in tissues and cells

We next studied the expression of SARS-CoV-2 associated genes and their expression profiles in different tissues and cell types. Only 6 genes (G3BP1, G3BP2, MARK3, PABPC1, PABPC4, and SRP54) overlapped between SARS-CoV-2 associated genes proposed by Gordon et al. and multiple CoVs associated genes proposed by Zhou et al. (Figure 2a and Supplementary Figure 2) (Gordon et al. 2020; Zhou et al. 2020). Enrichment analysis revealed that the overlapping genes belong to the ribonucleoprotein complex (GO:1990904) and stress granule assembly (GO:0010494) GO terms. However, due to the stark differences between the gene lists of SARS-CoV-2 and other coronavirus associated genes, we considered both these gene lists for further analysis.

Next, we calculated the gene score for both Gordon and Zhou gene lists in each cell type in each tissue from single cell RNA-Seq datasets. A positive gene score indicates that the genes in the list are expressed more than the background gene expression by the cell type. Interestingly, tissue resident immune cells such as dendritic cells and luminal macrophages from nasal and bronchi (Figure 2b, c), monocytes and T-cells from lung (Figure 2d), dendritic cells from oesophagus (Figure 2e), several lymphoid and myeloid cells from kidney (Figure 2f) exhibited significantly positive gene scores for SARS-CoV-2 and CoV associated genes. This potentially indicates that the host factors associated with coronaviruses are ubiquitously expressed by human immune cells (see Figure 2 and Supplementary Figure S3 for all single cell RNA-Seq datasets).

### 3.3 Circulating Immune cells as potential targets of SARS-CoV-2

To test if immune cells can potentially act as targets of SARS-CoV-2, we studied ACE2 and TMPRSS2 expression in circulating immune cells using bulk transcriptomics data. We found that ACE2 and TMPRSS2 are not expressed in any circulating immune cells (Figure 3a and b). We next performed machine learning based gene enrichment analysis for CoV and SARS-CoV-2 associated genes in bulk immune cell transcriptomics datasets using DIME (see methods). DIME is a tool specifically built for identifying key immune cell types and their corresponding key genes from a user defined gene list (Devaprasad, Radstake, and Pandit 2019). We found that several CoV associated genes extracted from Zhou et al. were highly expressed in T-cells and several of these genes were expressed in nearly all immune cell types (Figure 3c; see colors for score rank 1). The genes that were highly expressed in all immune cells included EEF1A1, PABPC1, RPL19, HNRNPA2B1, etc., and these genes correspond to pathways related to mRNA translation (Supplementary Figure 5a). The NSP-1 protein of CoV (Lokugamage et al. 2012) and SARS-CoV-2 (Thoms et al. 2020) has been shown to be involved in immune evasion by shutting down RNA translation. The exact interaction of SARS-CoV-2 and the genes involved in RNA translation in T-cells and other immune cells is unknown and needs to be further elucidated to see if there is a link between this interaction and the lymphopenia often seen in COVID-19 patients (Huang and Pranata 2020). Similarly, we found that macrophages and T-cells were enriched in SARS-CoV-2 associated genes extracted from Gordon et al. (Figure 3d; see colors for score rank 1 and 2). The key genes enriched were found to be those associated with RAB signaling and neutrophil degranulation (Supplementary Figure 5b). Other viruses such as HIV have been shown to use RAB signaling related host vesicular transport for viral assembly (Spearman 2018). The exact involvement of RAB signaling in SARS-CoV-2 viral assembly is yet to be explored.

### 3.4 Peripheral Immune cells as potential targets of SARS-CoV-2

On the contrary, in multiple different single cell RNA-Seq datasets from peripheral tissues a fraction of different immune cell types expressed ACE2 and TMPRSS2 genes (Figure 1b-d and Supplementary Figure 1). Specifically, 0.42% of monocytes in lungs expressed ACE2 gene and this fraction is comparable to the other lung cells such as AT2, AT1 and endothelial cells (Figure 1d). Similarly, 0.45% to 0.54% of monocyte derived phagocytes (dendritic cells and macrophages) in kidney, 0.12% of macrophages in ileum, and 0.59% of dendritic cells in oesophagus expressed ACE2 gene (Figure 1e and f). Interestingly, lymphocytes in alveoli, pancreas, oesophagus, and spleen did not express ACE2 gene but a small fraction of T-cells in kidney (0.7% of CD8+ T-cells, 0.49% of CD4+ T-cells, 0.98% of NK cells, and 0.1% of B-cells), colon (0.03% CD8+ T-cells and 0.02% of memory B-cells), lung (0.51% of T-cells), ileum (0.12% of T-cells), and heart (0.16% of T/NK cells and 0.09% B-cells) expressed ACE2 gene (Supplementary Figure 1). Although the fraction of ACE2 expressing immune cells is very small, it is comparable to the fractions observed for some of the cell types known to be infected by SARS-CoV-2 such as AT2, AT1, epithelial, and endothelial cells. In summary, we found that immune cells in periphery (Figure 2) and circulation (Figure 3) are enriched for CoV and SARS-CoV-2 associated genes.

### 3.5 Protein expression of the key SARS-CoV-2 associated genes found in circulating immune cells and in tissues

We found the protein expression of all the key (top 25 percent genes of all clusters from DIME) SARS-CoV-2 associated genes to be high in all the circulating immune cells, Supplementary Figure 7a. The Pearson correlation estimate between the RNA-Seq and protein data (of circulating immune cells) for 5 cell types was found to be > 0.4 for all cell types except CD4 naive T cell (r=0.32). The correlations were found to be statistically significant (p-value < 0.05) for all the cell types (Supplementary Figure 7b). In the peripheral cells, we observed that all the key genes except the integrins were expressed ubiquitously across all the tissues except the brain, endometrium, kidney, liver, skeletal muscle, spleen, and thyroid glands (Supplementary Figure 8).

### 3.6 Enrichment of SARS-CoV-2 associated genes in the PBMCs of COVID-19 patients

We first explored the expression of SARS-CoV-2 associated genes in the PBMCs of the healthy controls and COVID-19 patients. Here, we used the published single cell RNA-Seq dataset from the covid19 cell atlas (Wilk et al. 2020; Ballestar et al. 2020). The Wilk et al., data comprised single cell RNA-Seq of PBMCs taken from 7 COVID-19 patients (confirmed by positive SARS-CoV-2 nasopharyngeal swab by RT-PCR) and 6 healthy controls. We then studied the expression of ACE2 and TMPRSS2 across pooled healthy controls and the different types of COVID-19 patients (Figure 4a-d). In the pooled healthy controls, only a small fraction of NK cells (0.021 %) expressed ACE2, whereas TMPRSS2 was expressed by several immune cells such as B-cells, CD14 monocytes, CD4 memory and naive T-cells, dendritic cells and NK cells (Figure 4a). In the pooled COVID-19 patients, ACE2 was expressed in several immune cells such as dendritic cells, CD14 monocytes, CD4 memory T-cells, CD4 naive T-cells, CD8 effector T-cells, etc; TMPRSS2 was expressed in most immune cells (Figure 4b). Interestingly, in the pooled COVID-19 patients, dendritic cells expressed ACE2 but not TMPRSS2. In the COVID-19 patients, that didn’t require ventilator, ACE2 was expressed in CD14 monocytes, CD4 memory and naive T-cells, and dendritic cells; TMPRSS2 was expressed in several immune cells (Figure 4c). However, in the COVID-19 patients that developed acute respiratory distress syndrome (ARDS) and required ventilator, ACE2 was expressed in CD14 monocytes, CD8 effector T-cells, dendritic cells, stem cells and eosinophils (Figure 4d).

**Figure 4:**
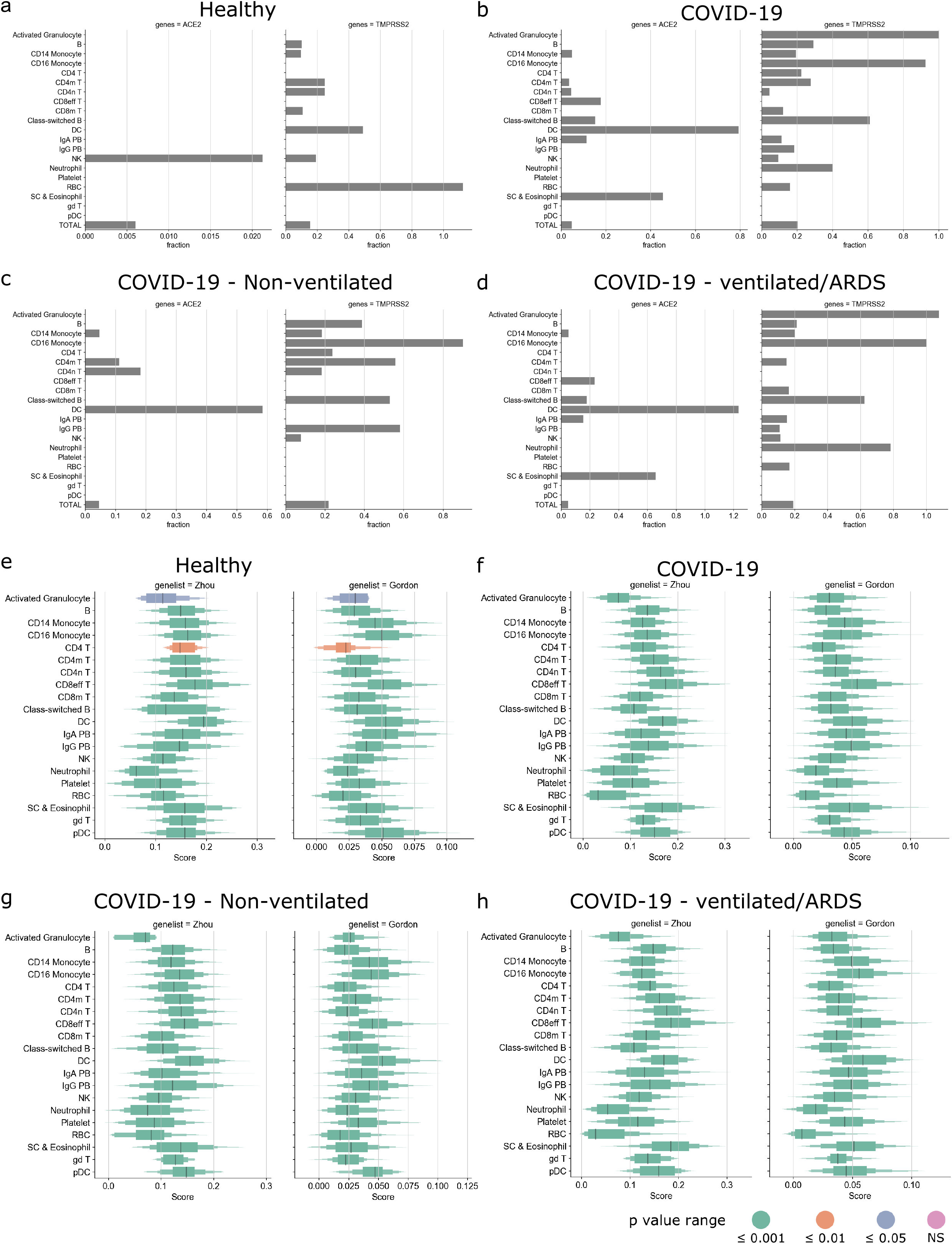
Enrichment of SARS-CoV-2 host factors in PBMCs of COVID-19 patients. Expression of ACE2 and TMPRSS2 in the pooled a. healthy controls, b. all COVID-19 patients, c. non-ventilated COVID-19 patients, and d. COVID-19 patients that developed ARDS. Boxplot showing distribution of gene score of Zhou and Gordon genes for e. healthy controls, f. all COVID-19 patients, g. non-ventilated COVID-19 patients, and h. COVID-19 patients that developed ARDS.

We then calculated the gene score for both Gordon and Zhou gene lists, for the pooled healthy controls and the different types of COVID-19 patients. The gene scores of Gordon and Zhou gene lists were significantly positive for all cells in the pooled healthy controls and the different types of COVID-19 patients (Figure 4e-h). Myeloid cells such as dendritic cells, CD14 and CD16 monocytes, plasmacytoid dendritic cells, and lymphoid cells such as CD8 effector T-cells and gamma delta T-cells had high gene scores across all types of patients. We then explored if the above patterns were discernable in the individual patient samples (Supplementary Figure 9a), we found that the fraction of ACE2 and TMPRSS2 expressing cells was heterogenous and did not show any patterns corresponding to the clinical outcome (Supplementary Figure 9b-i). However, upon inspection of the gene scores, we observed that, the gene scores for dendritic cells were consistently high across all patients (Supplementary Figure 9j-q), indicating that dendritic cells may play a key role in the infection dynamics.

## 4 Discussion

Understanding the pathophysiology and target cell populations is crucial for understanding the mechanism of action of SARS-CoV-2 and developing therapies for COVID-19. Here we used publicly available single cell RNA-Seq datasets from covid19 cell atlas (Sungnak et al. 2020; Wilk et al. 2020) to study which cells express ACE2 receptor and whether CoV (SARS-CoV-2 and other CoVs) associated genes are enriched in those cell types. We found that a tiny fraction of cells in tissues that are known to be targeted by SARS-CoV-2 express ACE2 gene, especially the lungs. In corroboration with Hou et al, we find a decreasing gradient in the fraction of ACE expressing cells from proximal to distal respiratory tract (Hou et al. 2020). A small fraction of ACE2 expressing immune cells were found in several tissues. Additionally, these immune cells in circulation and in the periphery are enriched in the SARS-CoV-2 associated genes.

Interestingly, circulating immune cells do not express ACE2 gene as shown from bulk transcriptome analysis. Multiple reasons can explain this finding. Firstly, we used bulk transcriptome data for circulating cells and hence we lose single cell resolution. However, we used bulk transcriptome data to study multiple conventional circulating cell types. Secondly, it could be that ACE2 expression is induced when the immune cells reside or reach different tissues and is not expressed when the immune cells are circulating. This is supported by the fact that multiple other cell types (such as enterocytes, endothelial cells, fibroblasts etc.) show differences in terms of ACE2 expression in different tissues. Lastly, the level of ACE2 expression in immune cells could be low and/or differs between cell and tissue types. We did not find this to be the case (Supplementary Figure 6), as ACE2 positive immune cells typically expressed ACE2 gene at a level that was comparable to the other SARS-CoV-2 target cells found in the corresponding tissues.

Additionally, we used publicly available single cell RNA-Seq dataset of PBMCs from COVID-19 positive patients to explore the expression of SARS-CoV-2 associated host factors (Figure 4 and supplementary Figure 9). We found that only a tiny fraction of cells expressed ACE2 and TMPRSS2. Interestingly, the fraction of ACE2 and TMPRSS2 expressing cells were found to be heterogenous across the different patients and was not representative of severity of disease or clinical outcome. However, on inspection of the SARS-CoV-2 associated host factors we found that several immune cells (such as dendritic cells, plasmacytoid dendritic cells, monocytes, T-cells, etc) were enriched similarly across the different COVID-19 patients, indicating that ACE2 and TMPRSS2 may not be sufficient to study SARS-CoV-2 infection.

Alternatively, some studies have indicated that SARS-CoV-2 might infect using additional receptors and co-receptors (Singh, Bansal, and Feschotte 2020; Sigrist, Bridge, and Le Mercier 2020). So we reperformed the entire analysis using entry factors compiled by Singh et al. and on all known integrins (Hynes 2002). We found that some integrins are expressed by some immune cells but most integrins are not expressed by most conventional immune cells in circulation and in tissues (Supplementary Figures 3-5). However, most of the immune cells found in the tissues are enriched in the entry factor list (including ANPEP, DPP4, FURIN) compiled by Singh et al. (Supplementary Figures 3 and 5c).

Given our computational findings, we propose that further studies should be performed to test if immune cells in periphery can express ACE2 and TMPRSS2 genes. Studying primary immune cells in circulation may not be sufficient to study ACE2 expression in immune cells. Recent preprint studies have shown that a few immune cell types (such as T-cells, B-cells, monocytes and macrophages) in tissues such as the lung, lymph nodes, and spleen are susceptible to SARS-CoV-2 infection (Pontelli et al. 2020; Boumaza et al. 2020; Feng et al. 2020). However, the immune cell types (including ACE2^-^ cells) susceptible to infection in all the other tissues remain uncharted. If even a small fraction of activated immune cells such as T-cells and dendritic cells can get directly infected by SARS-CoV-2, they might help the virus to migrate from one tissue to another. Thus, we propose host factors and immune cells that may play a role in viral replication, assembly and infection. Further research is needed to explore if SARS-CoV-2 can directly infect circulating and peripheral immune cells.

## 5 Conflict of Interest

The authors declare that the research was conducted in the absence of any commercial or financial relationships that could be construed as a potential conflict of interest.

## 6 Author Contributions

Conceptualization (AP), Formal Analysis (AD, AP), Methodology (AD, AP), Project Administration (AP), Supervision (AP), Visualization (AD), Writing original draft (AD, AP), and reviewing and editing manuscript (AD, AP).

## 7 Funding

Netherlands Organisation for Scientific Research (NWO) grant number: 016.Veni.178.027 (to AP).

## Notes

### Competing Interest Statement

The authors have declared no competing interest.

