## Supplementary Figure for "Enrichment of SARS-CoV-2 entry factors and interacting intracellular genes in peripheral immune cells"

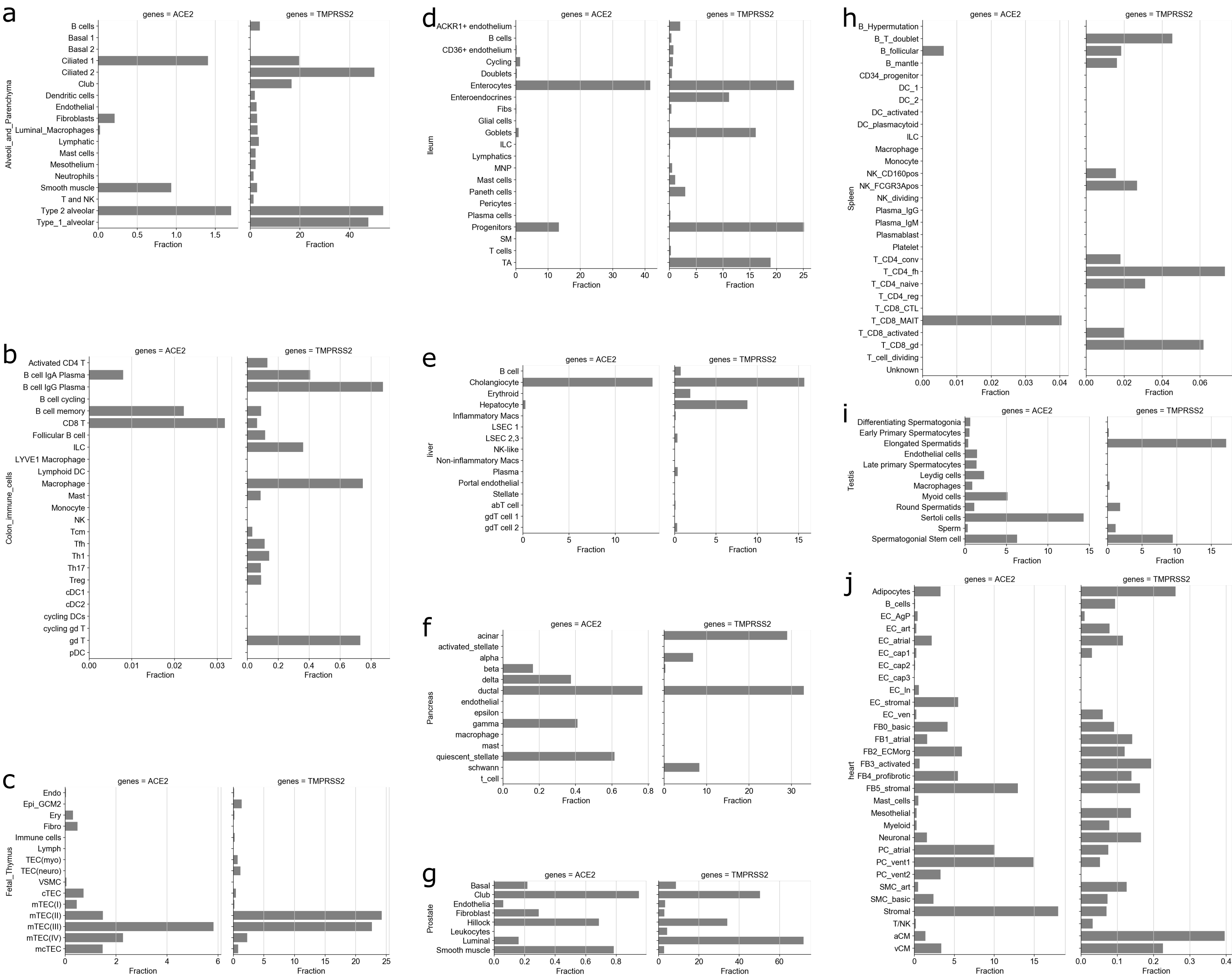

Supplementary Figure 1: Fraction plots of ACE2 and TMPRSS2 in different tissues. Fraction of cells (%) that express ACE2 and TMPRSS2.

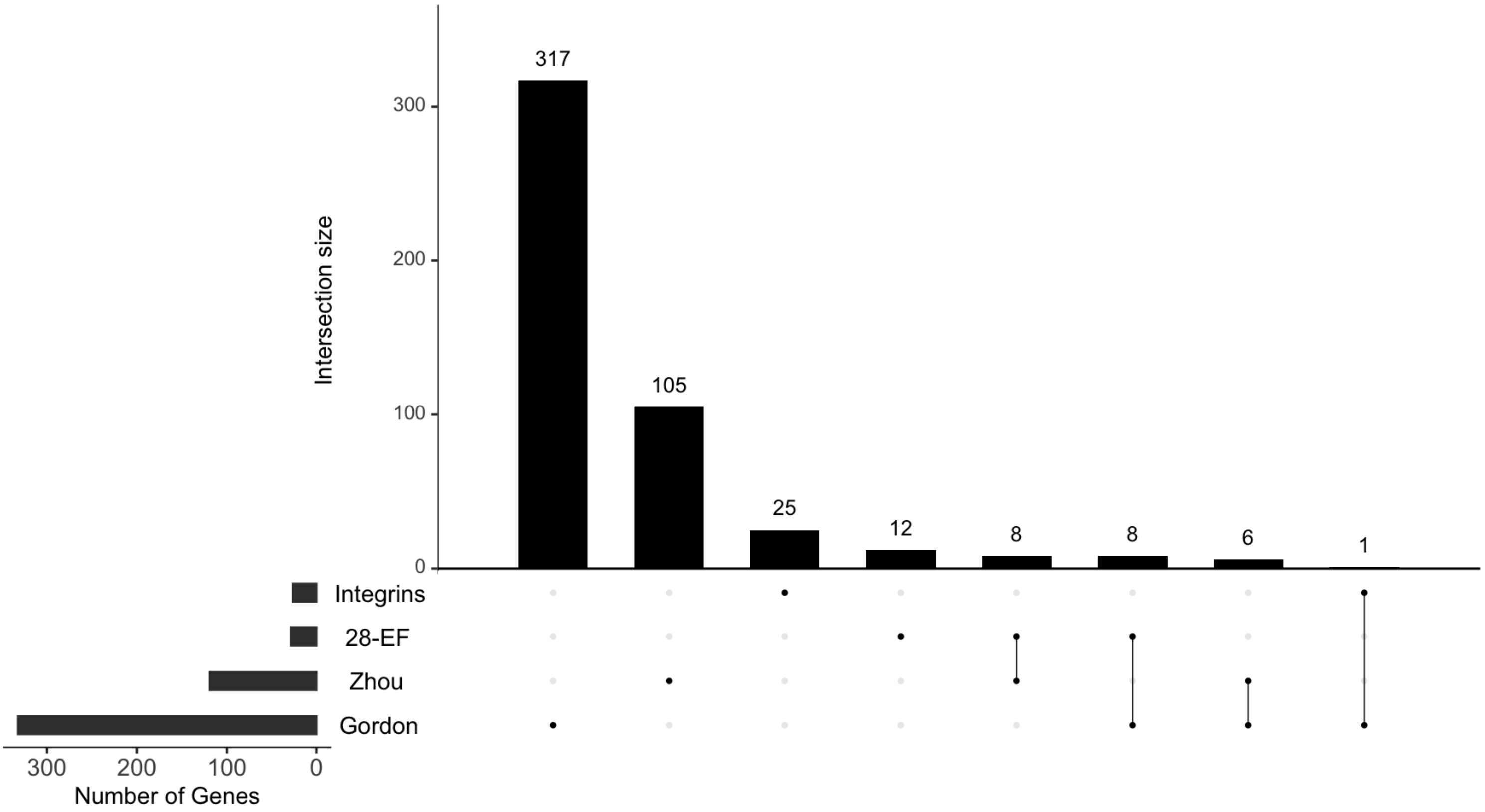


| | | | |
|---|---|---|---|
| ACE2 | AP2A2 | G3BP1 | ITGB1 |
| ANPEP | AP2M1 | G3BP2 | |
| CD209 | CHMP2A | MARK3 | |
| CLEC4G | RAB10 | PABPC1 | |
| CLEC4M | RAB14 | PABPC4 | |
| DPP4 | RAB1A | SRP54 | |
| TMPRSS2 | RHOA | | |
| ZCRB1 | TAPT1 | | |

Supplementary Figure 2: Intersection plot of different gene lists. Shows size of the gene lists in the bottom left panel, and overlapping size in the top panel. Genes overlapping between genelists are represented by vertical lines. Overlapping genes between the genelists are shown below the plot.

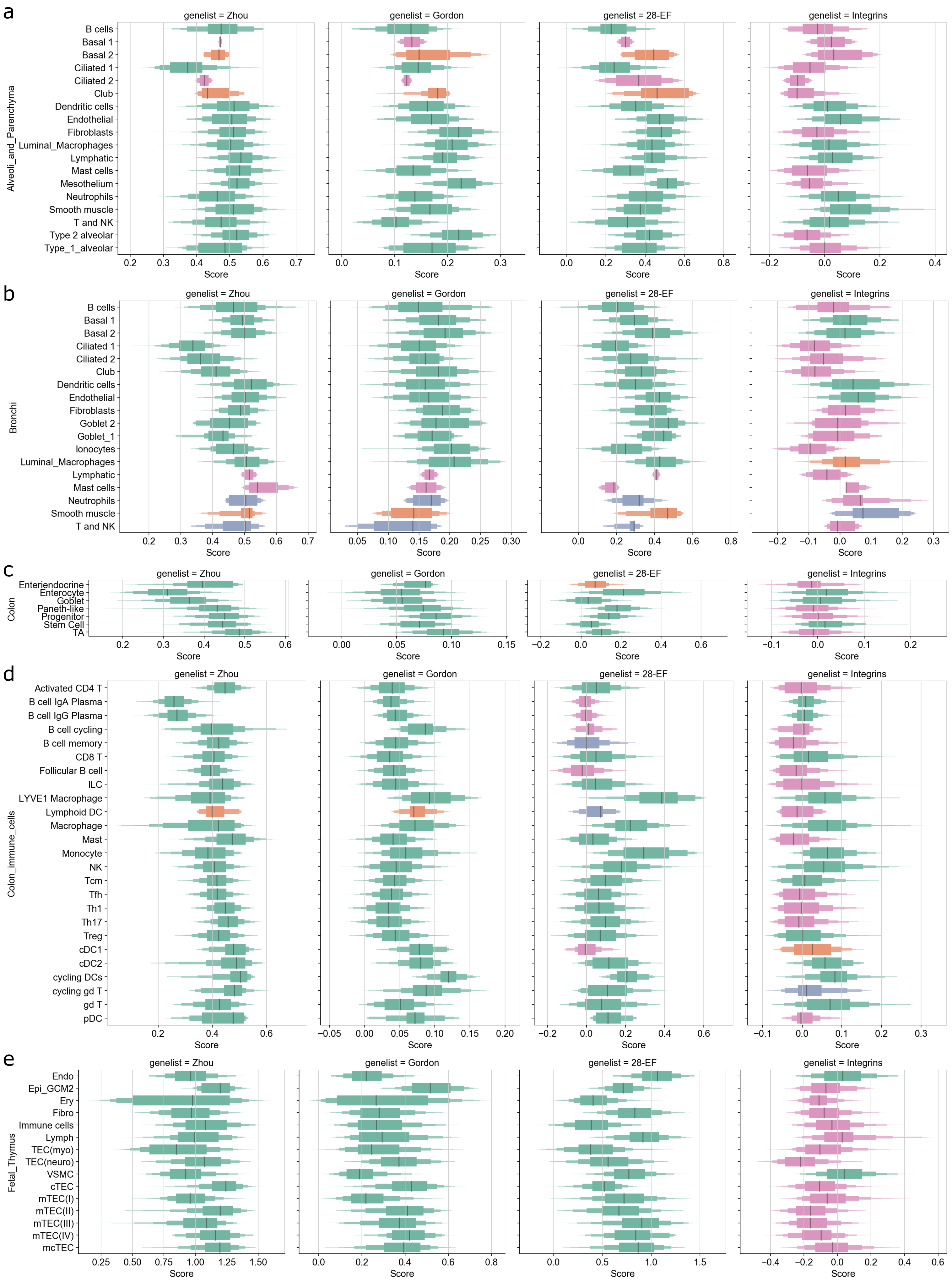

a
b
c
d
e
genelist = Zhou
genelist = Gordon
genelist = 28-EF
genelist = Integrins
Alveoli_and_Parenchyma
B cells
Basal 1
Basal 2
Ciliated 1
Ciliated 2
Club
Dendritic cells
Endothelial
Fibroblasts
Luminal_Macrophages
Lymphatic
Mast cells
Mesothelium
Neutrophils
Smooth muscle
T and NK
Type 2 alveolar
Type_1_alveolar
Bronchi
Goblet 2
Goblet_1
Ionocytes
Colon
Enteriendocrine
Enterocyte
Goblet
Paneth-like
Progenitor
Stem Cell
TA
Colon_immune_cells
Activated CD4 T
B cell IgA Plasma
B cell IgG Plasma
B cell cycling
B cell memory
CD8 T
Follicular B cell
ILC
LYVE1 Macrophage
Lymphoid DC
Macrophage
Mast
Monocyte
NK
Tcm
Tfh
Th1
Th17
Treg
cDC1
cDC2
cycling DCs
cycling gd T
gd T
pDC
Fetal_Thymus
Endo
Epi_GCM2
Ery
Fibro
Immune cells
Lymph
TEC(myo)
TEC(neuro)
VSMC
cTEC
mTEC(I)
mTEC(II)
mTEC(III)
mTEC(IV)
mcTEC
Score

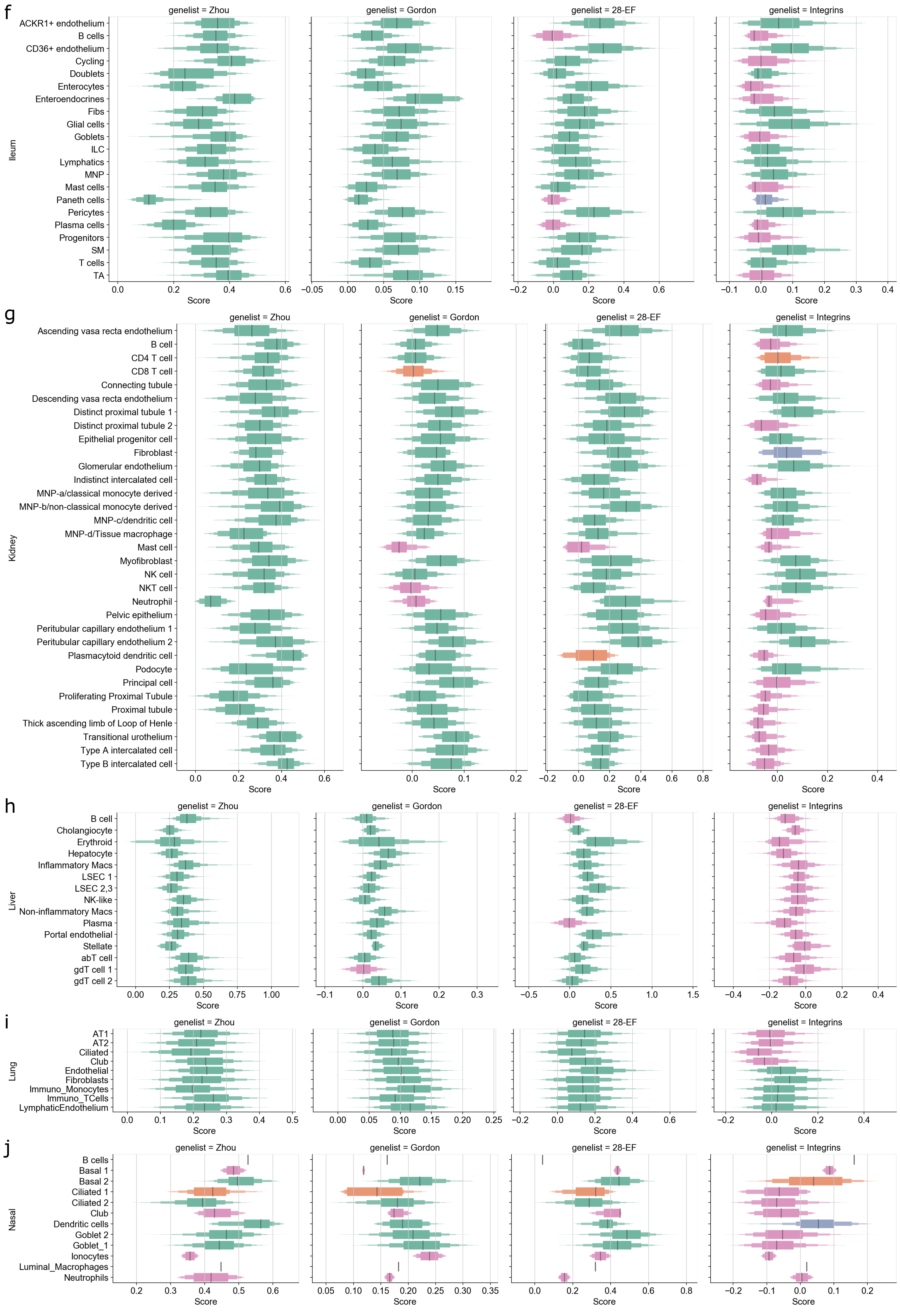
f
genelist = Zhou
genelist = Gordon
genelist = 28-EF
genelist = Integrins
Ileum
ACKR1+ endothelium
B cells
CD36+ endothelium
Cycling
Doublets
Enterocytes
Enteroendocrines
Fibs
Glial cells
Goblets
ILC
Lymphatics
MNP
Mast cells
Paneth cells
Pericytes
Plasma cells
Progenitors
SM
T cells
TA
Score
g
genelist = Zhou
genelist = Gordon
genelist = 28-EF
genelist = Integrins
Kidney
Ascending vasa recta endothelium
B cell
CD4 T cell
CD8 T cell
Connecting tubule
Descending vasa recta endothelium
Distinct proximal tubule 1
Distinct proximal tubule 2
Epithelial progenitor cell
Fibroblast
Glomerular endothelium
Indistinct intercalated cell
MNP-a/classical monocyte derived
MNP-b/non-classical monocyte derived
MNP-c/dendritic cell
MNP-d/Tissue macrophage
Mast cell
Myofibroblast
NK cell
NKT cell
Neutrophil
Pelvic epithelium
Peritubular capillary endothelium 1
Peritubular capillary endothelium 2
Plasmacytoid dendritic cell
Podocyte
Principal cell
Proliferating Proximal Tubule
Proximal tubule
Thick ascending limb of Loop of Henle
Transitional urothelium
Type A intercalated cell
Type B intercalated cell
Score
h
genelist = Zhou
genelist = Gordon
genelist = 28-EF
genelist = Integrins
Liver
B cell
Cholangiocyte
Erythroid
Hepatocyte
Inflammatory Macs
LSEC 1
LSEC 2,3
NK-like
Non-inflammatory Macs
Plasma
Portal endothelial
Stellate
abT cell
gdT cell 1
gdT cell 2
Score
i
genelist = Zhou
genelist = Gordon
genelist = 28-EF
genelist = Integrins
Lung
AT1
AT2
Ciliated
Club
Endothelial
Fibroblasts
Immuno_Monocytes
Immuno_TCells
LymphaticEndothelium
Score
j
genelist = Zhou
genelist = Gordon
genelist = 28-EF
genelist = Integrins
Nasal
B cells
Basal 1
Basal 2
Ciliated 1
Ciliated 2
Club
Dendritic cells
Goblet 2
Goblet_1
Ionocytes
Luminal_Macrophages
Neutrophils
Score

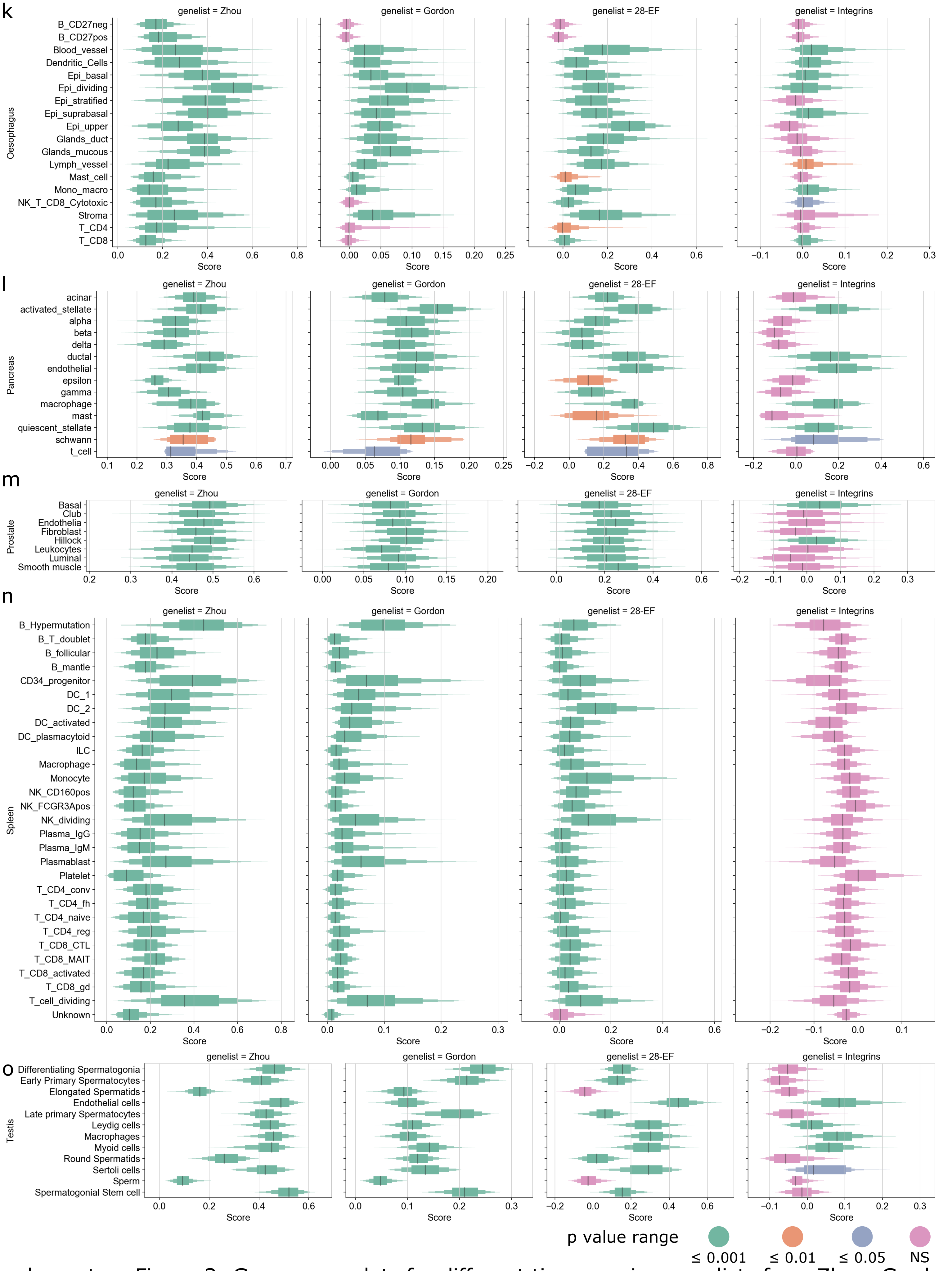


Supplementary Figure 3: Gene score plots for different tissues using genelists from Zhou, Gordon, 28-EF and Integrins. Gene score was calculated using tl.score_genes function from the scanpy suite. Positive gene score shows the given genes are expressed more than the background genes (all other genes). P value was calculated using the non parametric Wilcoxon test. The color of the boxen represent the p value ranges. NS is non significant, i.e, p value > 0.05.

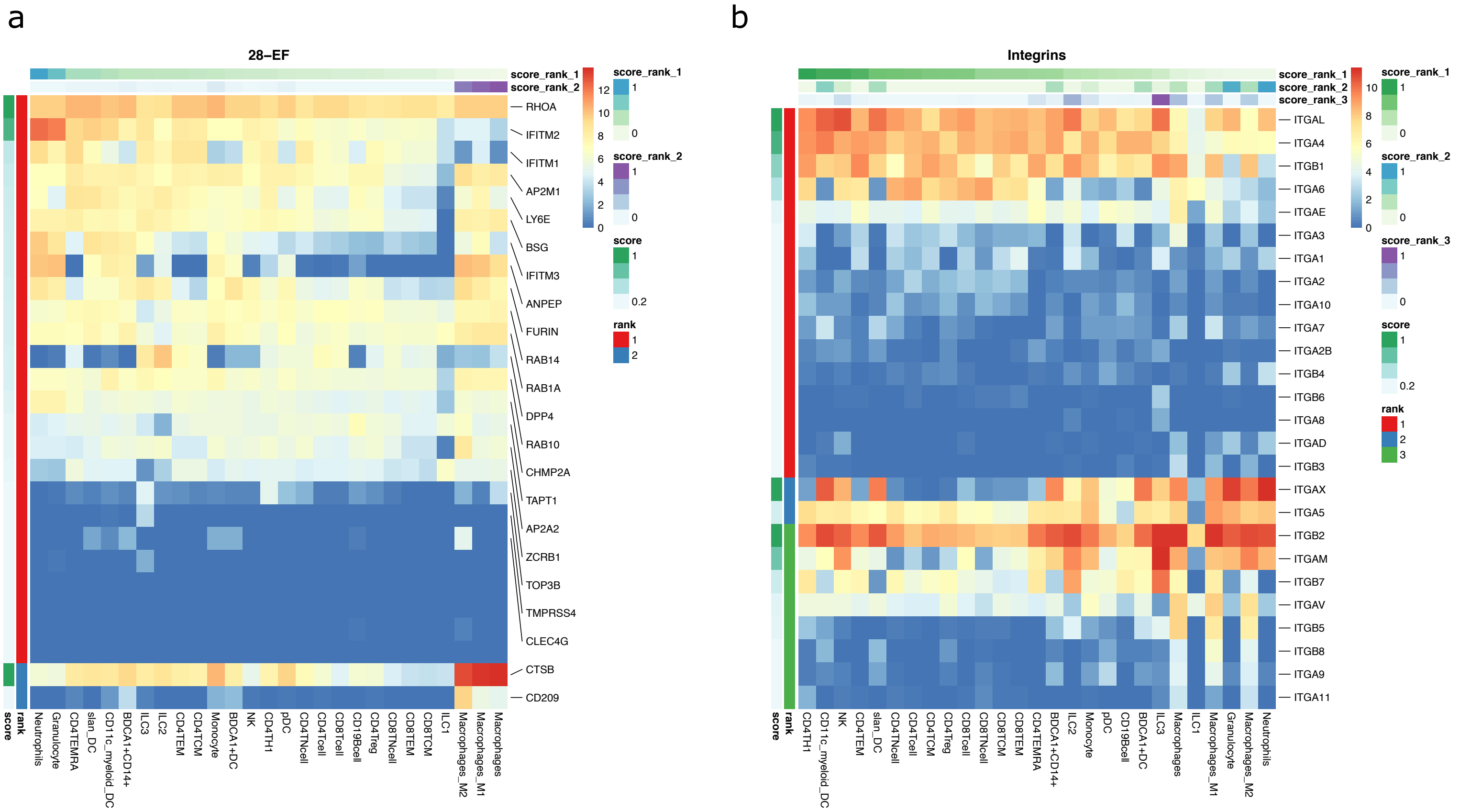


Supplementary Figure 4: DIME Heatmap showing ranked enrichment of a. 28-EF (Singh et al.), b. Integrins. The ranks depict the clusters as identified by the DIME (see methods). Top 20 genes (if present) for each rank are shown. The cells are ordered based on the score of the rank 1 (Top weighted rank). Expression values are in log2(cpm+1).

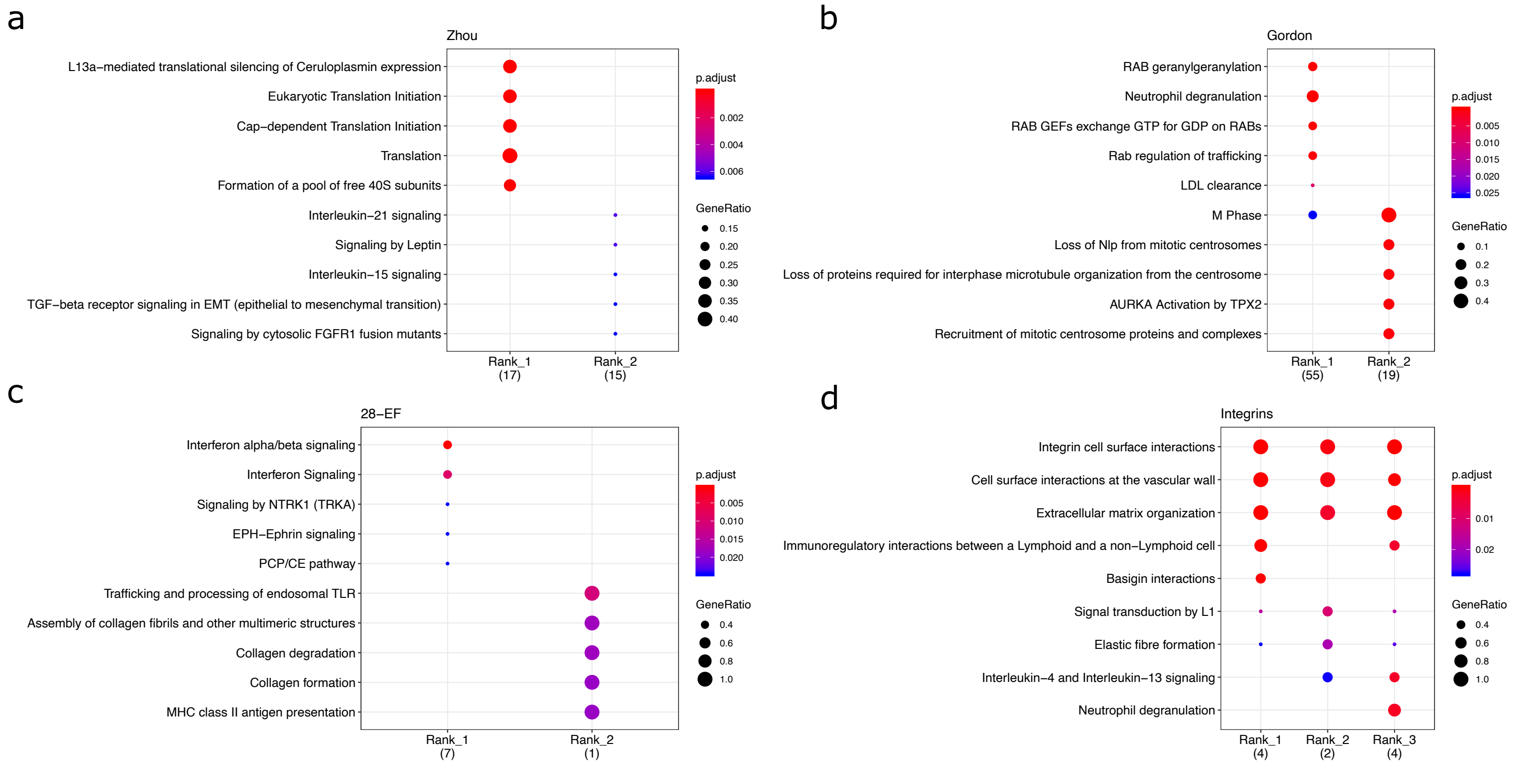


Supplementary Figure 5: Pathway analysis (Reactome) plots for the top 25 percent genes of each rank as calculated by DIME (and as depicted in Supplementary Figure 4) for the a. Zhou, b. Gordon, c. 28-EF and d. Integrin gene lists.

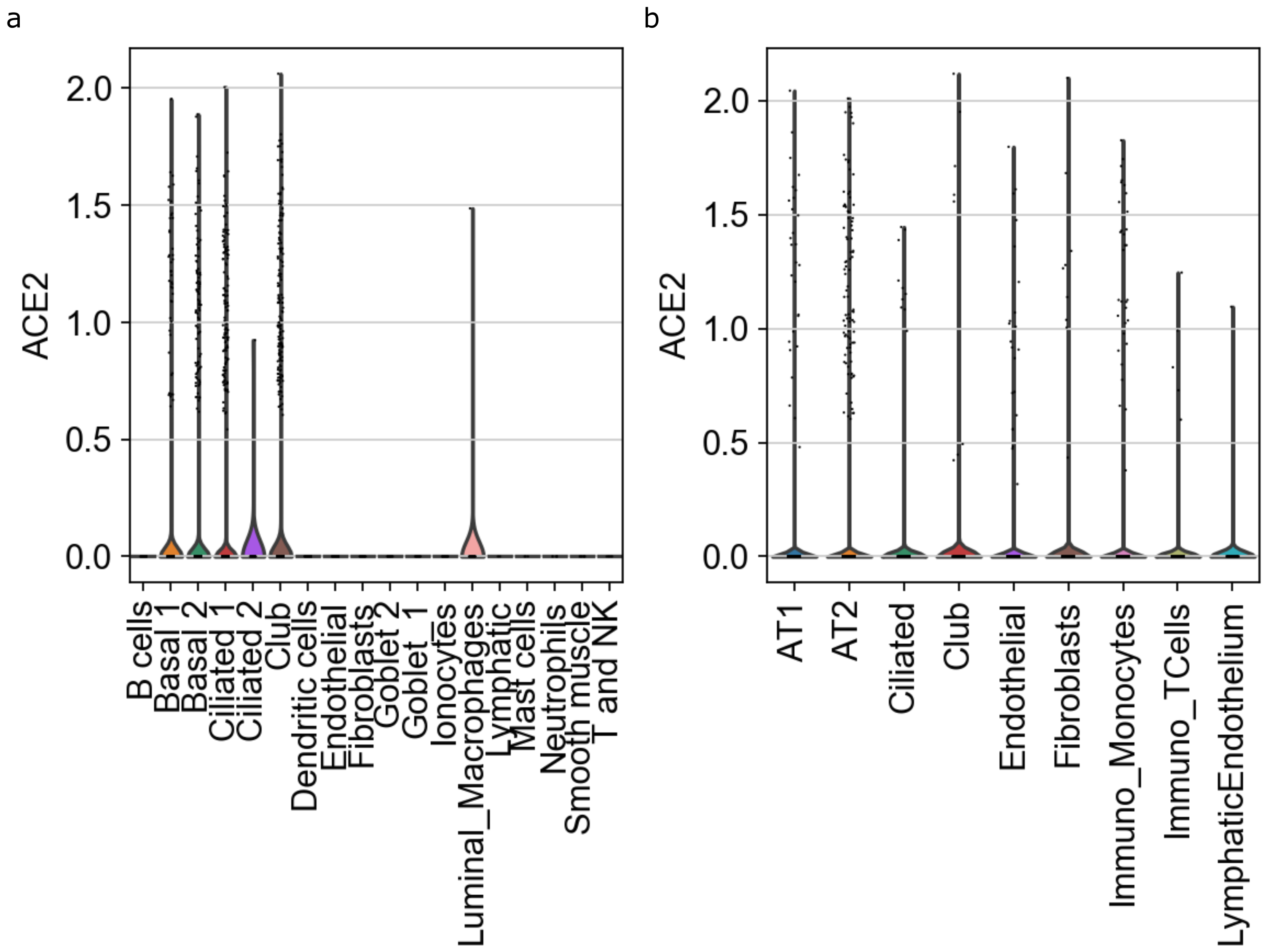


Supplementary Figure 6: ACE2 receptor gene expression of different cells across a. Bronchi and b. Lung

a

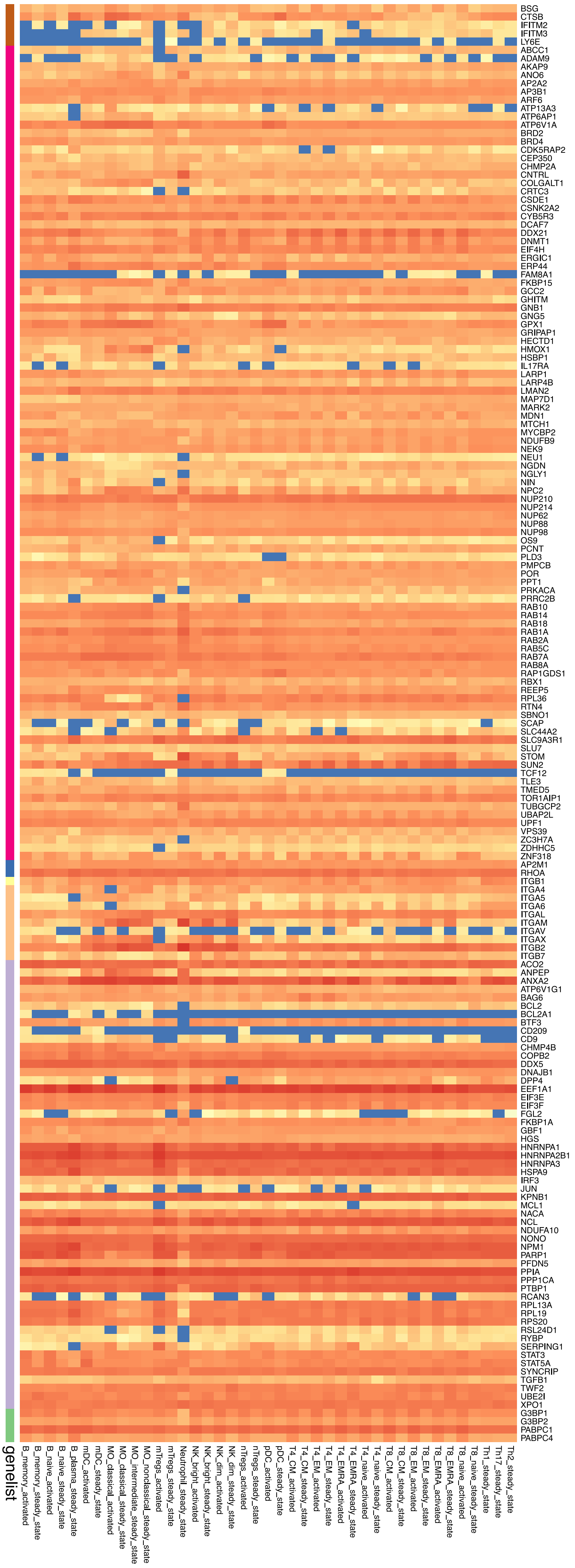


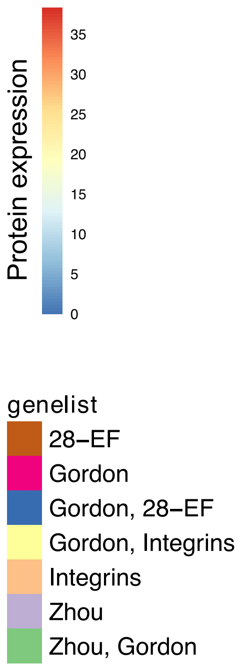


b

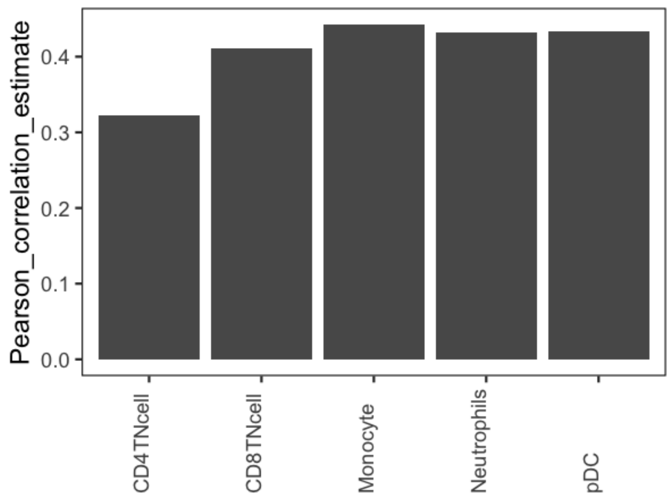


Supplementary Figure 7: a. Heatmap of protein expression of circulating immune cells from the immprot database. Shown here are protein expression of the top 25 percent genes (all clusters) of each genelist as identified by the DIME analysis (from Figure 3 and Supplementary Figure 4). Each row represents protein from the genelist (represented by color) and column is the median representative of the cell type. b. Pearson correlation estimate between RNA-Seq and immprot dataset shown for the 5 cell types. Pearson correlation p-value was found to be ≤ 0.05 for the 5 cell types.

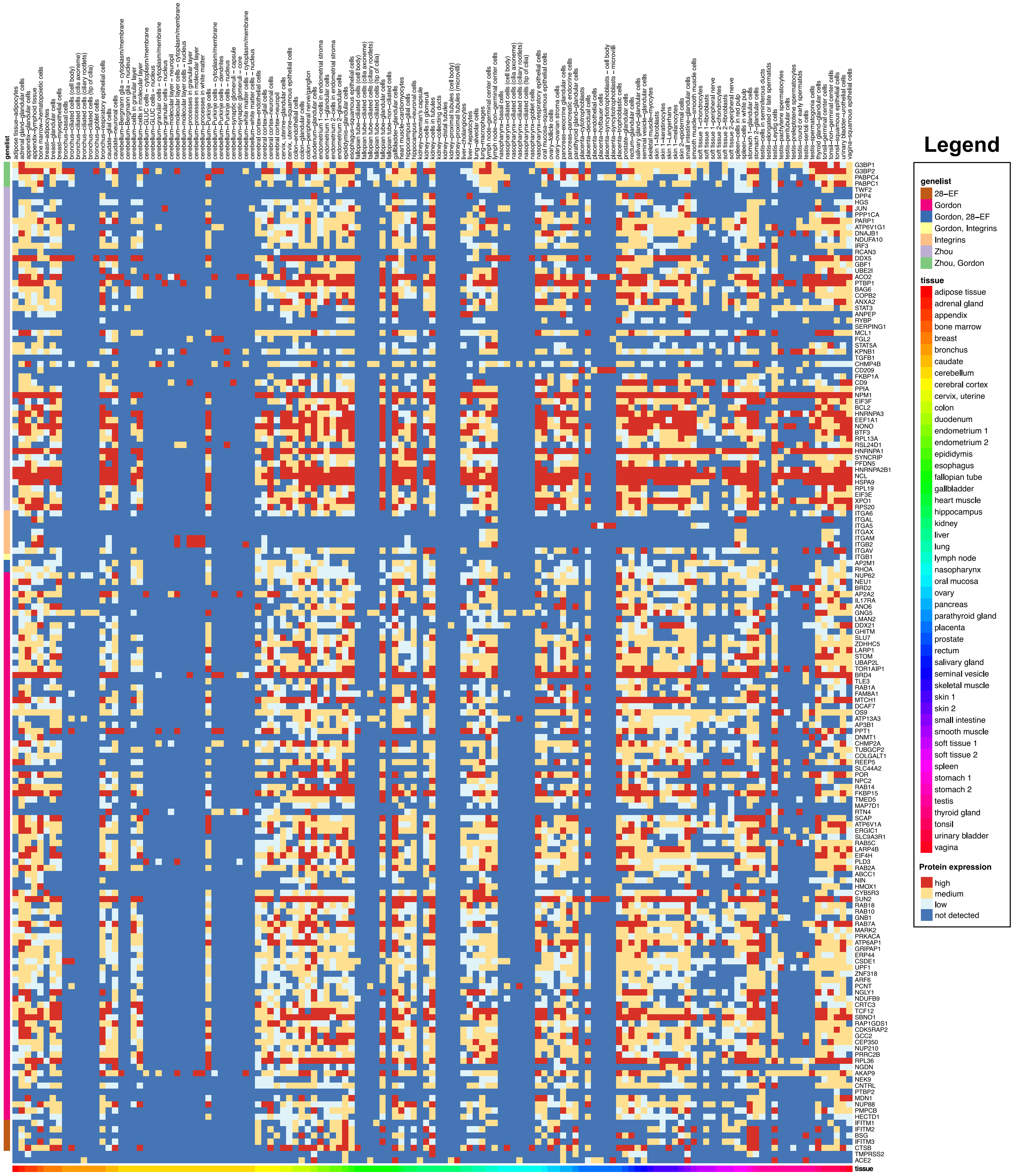

Supplementary Figure 8: Heatmap of protein expression of peripheral cells from 49 tissues taken from the Human Protein Atlas (HPA). Shown here are protein expression of the top 25 percent genes (all clusters) in each genelist as identified by the DIME analysis (from Figure 3 and Supplementary Figure 4). Each row represents protein from the genelist and the column represents cell type, see legend.

a

| Patient | Ventilated/ARDS | Age | Sex | Admission level | Clinical outcome |
|---|---|---|---|---|---|
| C1A | No | 60-69 | M | ICU | Discharged to rehab on room air |
| C1B | Yes | | | | |
| C2 | No | 40-49 | M | ICU | Discharged home |
| C3 | Yes | 30-39 | M | ICU | Tracheostomy, Prolonged ICU and hospital course |
| C4 | Yes | 30-39 | M | ICU | Discharged home |
| C5 | No | 50-59 | M | ICU | Discharged home |
| C6 | Yes | >80 | M | ICU | Deceased |
| C7 | No | 20-29 | M | Floor | Discharged home |

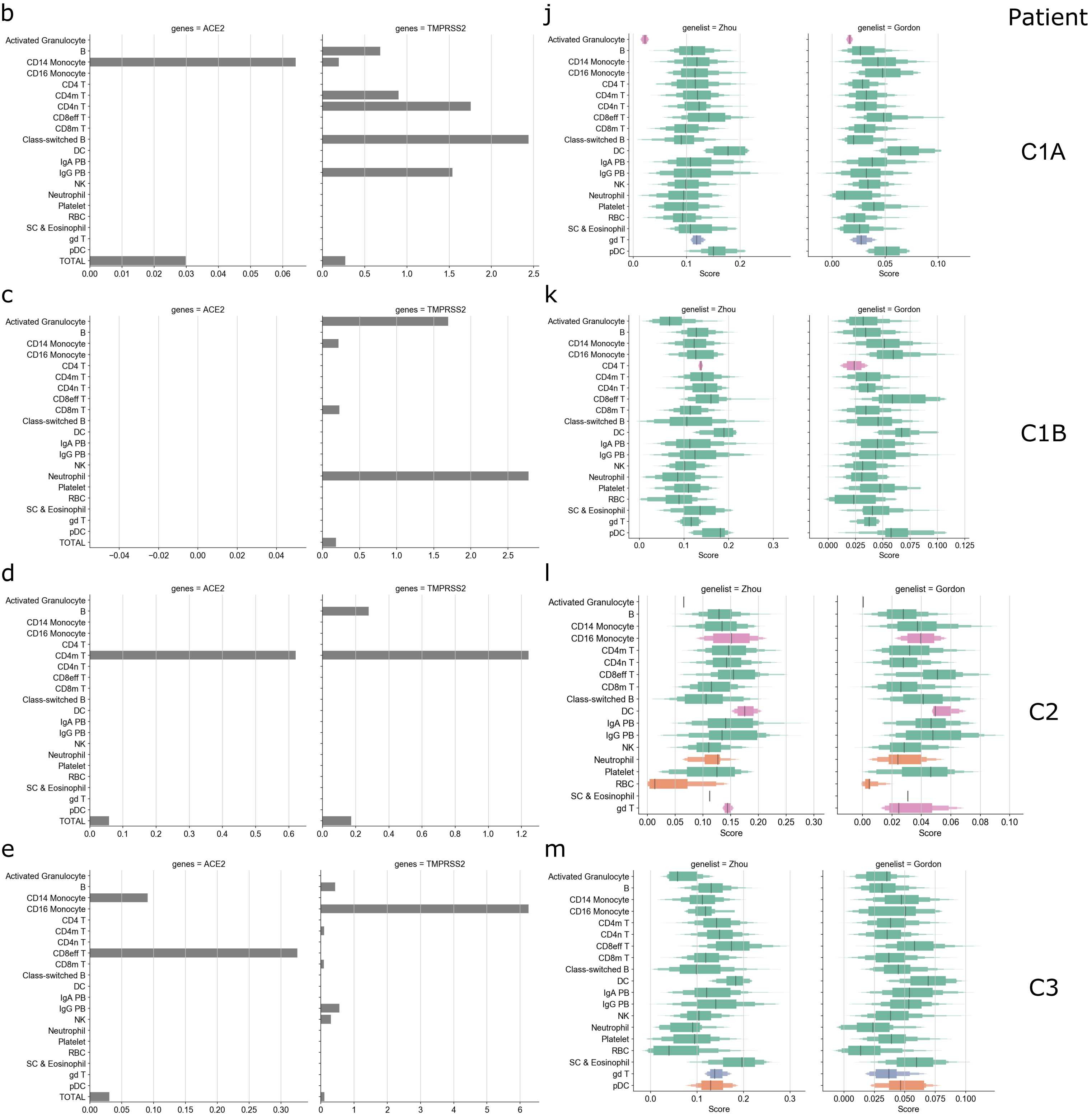

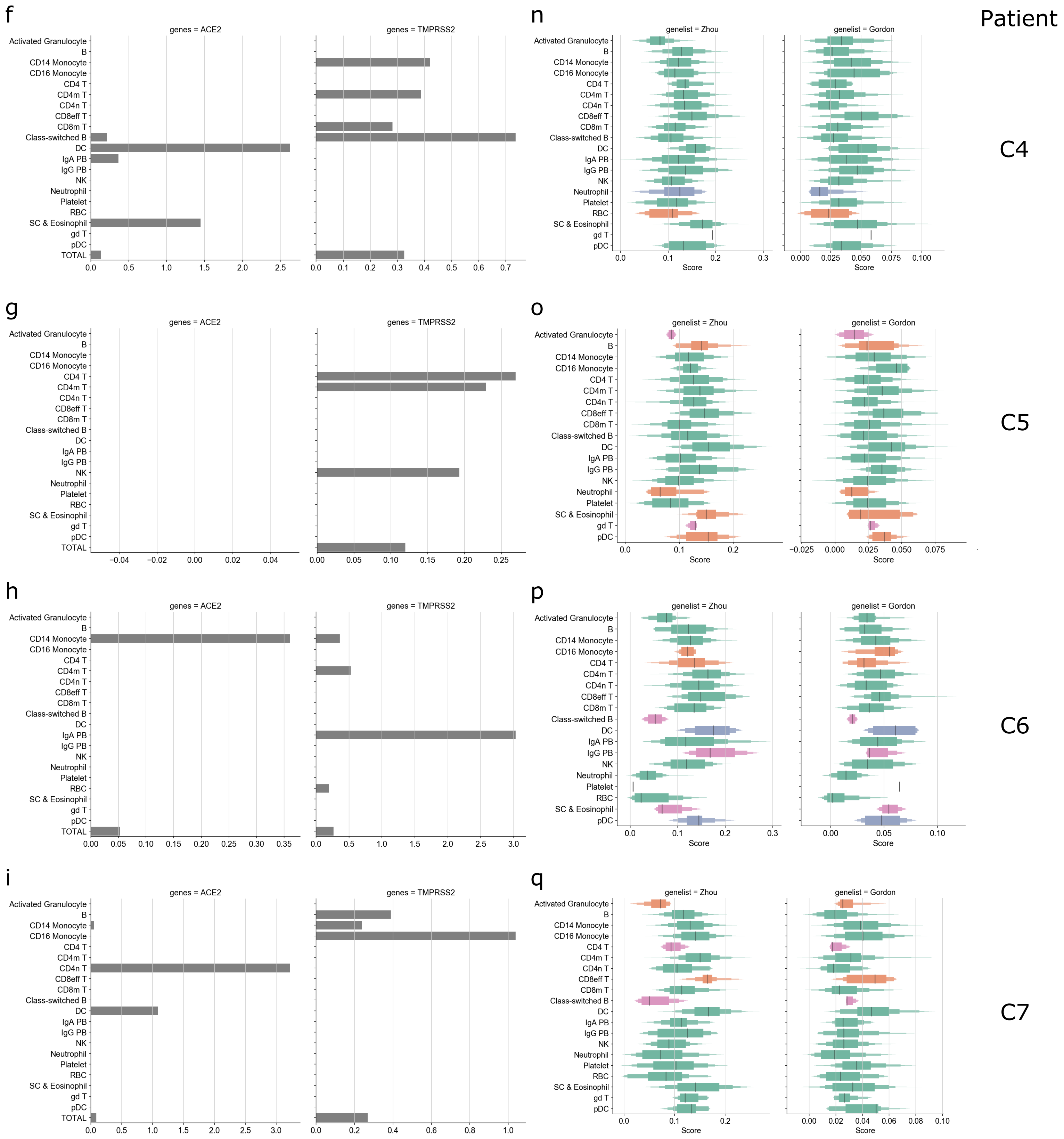


Supplementary Figure 9: a. Adapted from Wilk et al, the table represents the different COVID-19 patients and their clinical data. C1A and C1B represent the same patient, C1B was sample taken when the patient developed ARDS. b-i. represents the fraction of ACE2 and TMPRSS2 expressing cells across the different patient samples. j-q represents the Zhou and Gordon gene scores for the different patients. See Supplementary Figure 3 for p value legend.
